# Cellular integration with a subretinal honeycomb-shaped prosthesis

**DOI:** 10.1101/2022.12.15.520681

**Authors:** Mohajeet B. Bhuckory, Zhijie Chen, Bing-Yi Wang, Andrew Shin, Tiffany Huang, Ludwig Galambos, Efstathios Vounotrypidis, Keith Mathieson, Theodore Kamins, Daniel Palanker

**Affiliations:** Hansen Experimental Physics Laboratory, Stanford University, Stanford, CA, USA; Department of Ophthalmology, Stanford University, Stanford, CA, USA; Department of Electrical Engineering, Stanford University, Stanford, CA, USA; Department of Physics, Stanford University, Stanford, CA, USA; Department of Material Science, Stanford University, Stanford, CA, USA; Department of Physics, University of Strathclyde, Glasgow, Scotland, UK

## Abstract

In patients blinded by geographic atrophy, subretinal photovoltaic implant with 100µm pixels provided visual acuity closely matching the pixel pitch. However, such flat bipolar pixels cannot be scaled below 75µm, limiting the attainable visual acuity. This limitation can be overcome by shaping the electric field with 3-dimensional electrodes. In particular, elevating the return electrode on top of honeycomb-shaped vertical walls surrounding each pixel extends the electric field vertically and decouples its penetration into tissue from the pixel width. This approach relies on migration of the retinal cells into the honeycomb wells. Here, we demonstrate that the majority of the inner retinal neurons migrate into 25µm deep wells, leaving the third-order neurons, such as amacrine and ganglion cells, outside. This is important for selective stimulation of the second-order neurons to preserve the retinal signal processing in prosthetic vision. Comparable glial response to that with flat implants suggests that migration and separation of the retinal cells by the walls does not cause additional stress. Furthermore, retinal migration into the honeycombs does not negatively affect its electrical excitability.

## Introduction

Retinal degenerative diseases, such as age-related macular degeneration (AMD) and retinitis pigmentosa (RP), are among the leading causes of untreatable visual impairment. Despite the different pathophysiology, both diseases ultimately lead to loss of the photoreceptors, while leaving the inner retinal neurons relatively intact [1]–[3], albeit with some rewiring [4], [5]. Electrical stimulation of the remaining inner retinal neurons can elicit visual percepts, thereby enabling restoration of sight [6]–[8].

We developed an optoelectronic substitute for the lost photoreceptors: a wireless photovoltaic subretinal implant activated by light [9], [10]. Images of the visual scenes captured by a video camera are processed and projected by augmented-reality glasses onto a subretinal photodiode array using intense pulsed light. Photovoltaic pixels in the array convert this light into biphasic pulses of electric current, which stimulate the second-order neurons in the inner nuclear layer (INL) – primarily the bipolar cells (BC). To avoid perception of this light by the remaining photoreceptors in the peripheral region, we use a near-infrared (NIR, 880 nm) wavelength.

This approach offers multiple advantages: (1) thousands of pixels in the implant can be activated simultaneously and independently; (2) a lack of wires enables reliable encapsulation of the implant and greatly simplifies the surgical procedure; (3) beside autofocusing, an external camera allows operation over a wide range of ambient illumination and provides adjustable image processing optimized for the dynamic range of the implant; (4) light-sensitive pixels maintain the natural link between eye movements and image perception; (5) network-mediated retinal stimulation retains many features of the natural signal processing, including antagonistic center-surround [11], flicker fusion at high frequencies and nonlinear summation of the RGC subunits [9], amongst others.

This approach has been employed clinically, where PRIMA implants (Pixium Vision SA, Paris, France) with 100µm pixels, were implanted in AMD patients. These patients perceived monochromatic formed vision in the previous scotomata, with a prosthetic visual acuity closely matching the pixel size: 1.17±0.13 pixels, corresponding to the Snellen range of 20/438–20/550 (100 µm pixel corresponds to 20/420 acuity) [12], [13]. Even more remarkable is that the prosthetic central vision in AMD patients is perceived simultaneously with the remaining natural peripheral vision [13].

However, for a wider adoption of this approach by AMD patients, prosthetic acuity should significantly exceed that of their remaining peripheral vision, which is typically no worse than 20/400. The Nyquist sampling limit for an acuity of 20/200 corresponds to 50 µm pixels, with 20/100 equating to 25 µm. As with natural vision, prosthetic acuity is limited not only by the spatial resolution of the stimulation patterns (i.e. pixel size and the field spread), but also by their contrast, which is affected by crosstalk between the neighboring electrodes [14]. Lateral spread of electric field can be confined by the local return electrodes in each pixel, as in the PRIMA implant (Figure 1a), but scaling down such pixels is difficult because the penetration depth of the electric field in tissue is constrained by the distance between the active and the return electrode, which is about half of the pixel radius [15]. As a result, the stimulation threshold in such geometry rapidly increases with a decreasing pixel size, and exceeds the safe charge injection limit for pixels below 75 µm in human retina [16] and 55 µm in rats [17], even with one of the best electrode materials – sputtered iridium oxide films (SIROF) [15].

**Figure 1.**
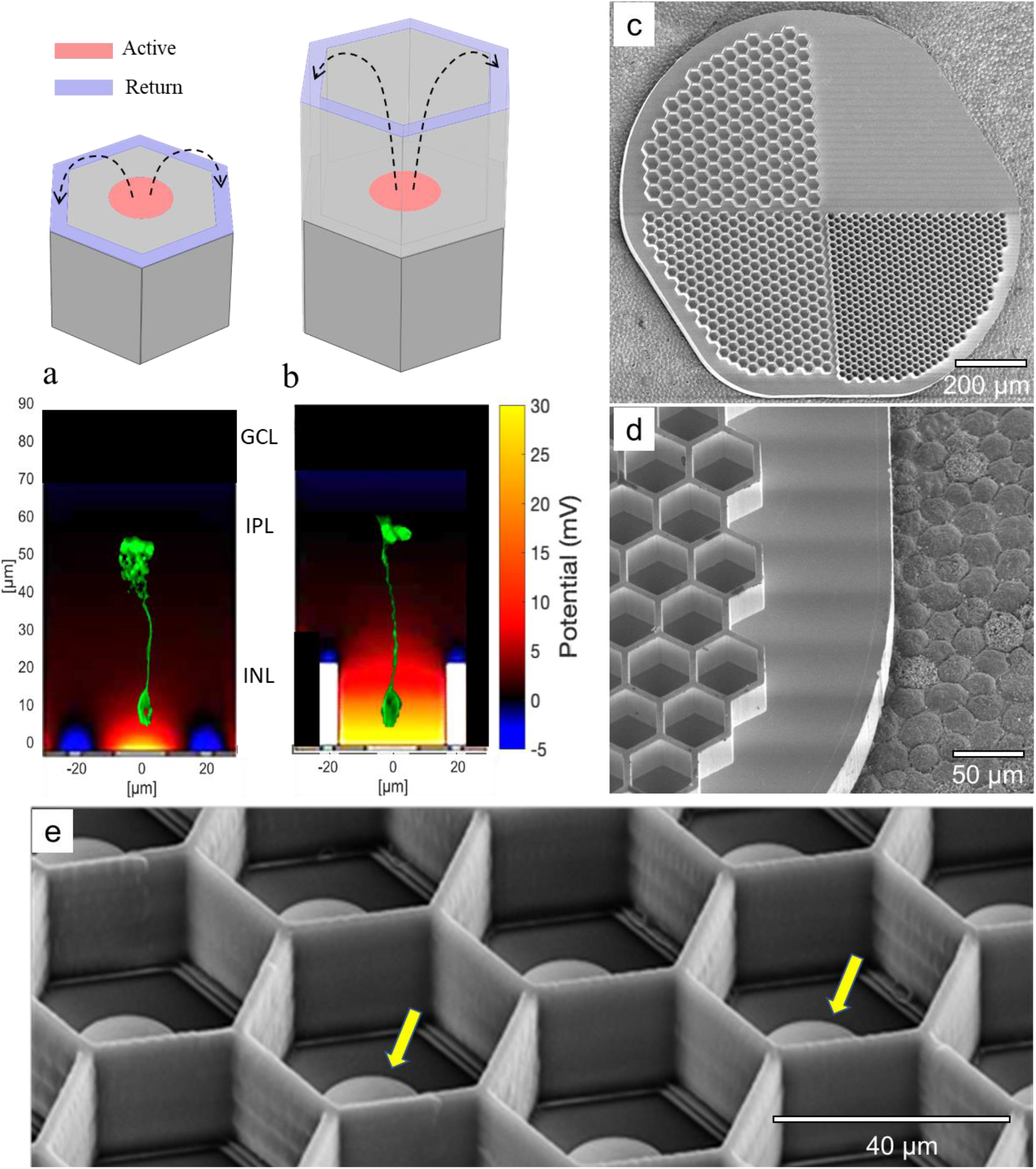
Photovoltaic pixels of 40 *µm* in width with different return electrode geometries. (a) A flat bipolar pixel containing the central active and a circumferential return electrode. (b) A honeycomb-shaped pixel with elevated return electrode. Bottom panels show the corresponding simulated electric potential (current = 68nA). A PKCα labelled bipolar cell with its axonal terminals in the middle of IPL in the left panel illustrates its position and size with respect to the field penetration. SEM of the honeycomb devices: (c) Passive implants with 4 quadrants: flat, 20, 30 and 40 µm wells. (d) Higher magnification of the passive 40 µm wells imaged on porcine RPE for size illustration. (e) 40 µm photovoltaic pixels with polymer honeycomb walls. Yellow arrows point at the active electrodes.

To overcome this limitation, we proposed a 3-dimentional (3-D) “honeycomb” configuration of the implant, with the active electrodes at the bottom and return electrodes at the top of vertical walls [15] (Fig. 1b). In this configuration, the electric field in the cavities is oriented nearly vertically, parallel to the direction of the bipolar cells, thereby reducing the stimulation threshold. Decoupling the electric field penetration depth (defined by the cavity height) from the pixel width enables scaling the pixel size down to cellular dimensions. Furthermore, confinement of the electric field within cavities limits the cross talk from neighboring pixels.

The honeycomb approach is based on retinal migration: within days after the implantation, inner retinal neurons migrate into open spaces in a subretinal implant [18], [19]. For the network-mediated retinal stimulation, the second-order neurons should be activated below the threshold of the direct stimulation of the third-order retinal neurons. Therefore, the wall height must accommodate the bipolar cells within the cavities, while leaving the amacrine (AC) and ganglion cells (RGCs) outside.

Here, we present a detailed anatomical characterization of the retinal migration in Royal College Surgeon (RCS) rats after subretinal implantation of the honeycomb arrays with 20, 30 and 40 µm pixels. We also assessed the potential effect of this migration on retinal excitability by measuring the visually evoked potential (VEP). Our results suggest that retinal prostheses with such 3-D structures enable the retinal stimulation with pixels down to 20 µm in size, corresponding to a visual acuity of 20/80, which would significantly help many patients impaired by atrophic AMD.

## Methods

### Honeycomb implants

The Inner Nuclear Layer (INL) in RCS rats is about 40-50µm thick and is comprised of 4-5 layers of nuclei. Assuming that amacrine cells (AC) nuclei are located in the top third of the INL, we made the walls 25 µm in height. Passive honeycomb implants for anatomical studies were fabricated from crystalline silicon using two mask layers to generate patterns for deep silicon etching, as previously described [15]. A hexamethyldisilazane (HMDS) primed wafer was spin-coated with 2 µm of negative photoresist (AZ5214-IR), which was then exposed to UV light through a pattered photomask. 25 µm deep cavities were then etched in the unprotected regions using a Bosch etch process. After the honeycomb-defining resist was removed, photoresist (7.5% SPR 220-7, 68% MEK, and 24.5% PGMEA) was spray-coated over the wafer to a thickness of 30 μm and exposed to define the releasing trenches around the 1 mm wide arrays, also using a Bosch process. The wafer was then spray-coated with a protective 60 µm thick photoresist for the backside grinding from 500 to 50 µm in thickness from the base of the honeycombs. Subsequent etching of the remaining excess silicon in XeF_2_ gas completed the release of the implants. Each implant had four quadrants with hexagonal honeycomb patterns of 40, 30 and 20 µm in width and having 25 µm high walls of 4, 3, and 2 µm thicknesses, respectively. The fourth quadrant served as a flat control, shown in Fig. 1c & 1d. Arrays were sputter-coated with 200 nm of gold to prevent dissolution of the oxidized silicon (300 nm) in-vivo.

For the studies of retinal stimulation, 25 µm tall polymer walls were made with two-photon lithography (Nanoscribe GmbH & Co. KG, Karlsruhe, Germany) atop monopolar photovoltaic arrays [20]. These arrays are 1.5 mm in diameter, containing a 1.2 mm-wide hexagonal grid of either 821 pixels of 40 µm in size, or 2806 pixels of 20 µm. Each pixel has a vertical-junction photodiode with the anode connected to a disk coated with SIROF as the active electrode, and the cathode connected to a large annulus SIROF electrode as the global return in the periphery of the array [20]. The resulting active honeycomb implant with 40 µm pixels is shown in Fig. 1e.

### Animals and surgical procedures

All experimental procedures were approved by the Stanford Administrative Panel on Laboratory Animal Care (APLAC) and conducted in accordance with the institutional guidelines and conformed to the Statement for the Use of Animals in Ophthalmic and Vision research of the Association for Research in Vision and Ophthalmology (ARVO). Animal care and subsequent implantations were conducted as previously described [21] using rats with retinal degeneration from a Royal College of Surgeons (RCS) colony maintained at the Stanford Animal Facility. Total of N = 33 animals were implanted with passive honeycomb arrays and N = 10 animals with active honeycomb arrays after age of P180 to ensure complete degeneration of the photoreceptors. Animals were anesthetized with a mixture of ketamine (75 mg/kg) and xylazine (5 mg/kg) injected intraperitoneally. A 1.5 mm incision was made through the sclera and choroid 1 mm posterior to the limbus. The retina was detached with an injection of saline solution, and the implant was inserted into the subretinal space at least 3 mm away from the incision site. The conjunctiva was sutured with nylon 10-0, and topical antibiotic (bacitracin/polymyxin B) was applied on the eye postoperatively. Surgical success and retinal reattachment were verified using Optical Coherence Tomography (OCT) (HRA2-Spectralis; Heidelberg Engineering, Heidelberg, Germany).

### Retinal immunohistochemistry

For the retinal imaging and analysis, animals were euthanized 6-9 weeks post implantation using an intracardiac injection of Phenytoin/pentobarbital (Euthanasia Solution; VetOne, Boise, ID, USA). The eyes were enucleated and rinsed in phosphate buffered saline (PBS), anterior segment and lens were removed, the eye cup was cut to a 3 × 3 mm square centered around the implant, and fixed in 4% paraformaldehyde (PFA; EMS, PA, USA) for 12 hours at 4° C. Samples were permeabilized with 1% Triton X-100 (Sigma-Aldrich, CA, USA) in PBS for 3 hours at room temperature, followed by a blocking step in 10% bovine serum albumin (BSA) for 1 hour at room temperature, and a 48-hour incubation at room temperature with primary antibodies (Supplementary table 1) in 0.5% Triton X-100, 5% BSA in PBS. Samples were washed for 6 hours at room temperature in 0.1% Tween-20 in PBS (PBS-T), incubated for 48 hours at room temperature with secondary antibodies (Supplementary table.1) and counterstained with 4’, 6-Diamidin-2-phenylindol (DAPI) in PBS. After 6 hours of washing in PBS-T, the samples were mounted with Vectashield medium (H-1000; Vector Laboratories, Burlingame, CA, USA).

### Whole-mount retinal imaging and analysis

3-D imaging of the retinal whole mounts was performed using a Zeiss LSM 880 Confocal Inverted Microscope with Zeiss ZEN Black software. The implant surfaces were identified by reflection of a 514 nm laser with a neutral-density beam splitter allowing 80% transmission and 20% reflection. The images were acquired through the total thickness of the retina using a Z-stack, with the upper and lower bounds defined at the inner limiting membrane (ILM) and 10 μm below the base of the honeycomb wells, respectively. Stacks were acquired in the center of each honeycomb quadrant using a 40x oil-immersion objective with acquisition area >225×225 μm and 380 - 470 nm z-steps. The Zeiss z-stack correction module was used to account for dimmer light within the wells of the implants.

Confocal fluorescence datasets were processed using the FiJi distribution of ImageJ [22]. To correct for brightness variations at different Z positions in the stack within the wells and above the implant, we first maximized the contrast in the individual XY planes to ensure 0.3% channel saturation. The XY planes were de-speckled with the median filter and the background was suppressed using the rolling-ball algorithm [23]. The images then underwent cascades of gamma adjustments and min-max corrections to further suppress the background, depending on the noise level. Gaussian blurring was applied for nucleus staining channels to smoothen the brightness variations within individual cells. The implants were reconstructed by extruding the implant reflection toward the bottom (extraocular side) of the image stack.

To quantify the number of cells in the wells, the 3-D image of each honeycomb well was segmented into voxels based on the reflection channel using the Moore-Neighbor tracing algorithm implemented by the “bwboundaries” function in MATLAB 2021b (Mathworks, Inc., Natick, MA), while a control image stack (without an implant) was treated as one segment. Voxels brighter than 15% of the maximum intensity were considered positive, and in each segment, we defined three metrics as follows: (1) cell count – the manually counted total number of cell bodies or nuclei; (2) filling percentage – a fraction of positive voxels; (3) migration depth – the 95^th^ percentile of the depths for all positive voxels, counting from the top of the segment.

### Electrophysiology

For measurement of the visually evoked potential (VEP), each animal was implanted with three transcranial electrodes: one electrode above the visual cortex of each hemisphere (4 mm lateral from midline, 6 mm caudal to bregma), and a reference electrode in the somatosensory cortex (2 mm right of midline and 2 mm anterior to bregma).

For the pattern projection, following anesthesia and pupil dilation, viscoelastic gel was put between the cornea and a cover slip to cancel the corneal optical power and ensure good retinal visibility. The subretinal implant was illuminated using a customized projection system, including a near-infrared laser at 880 nm wavelength (MF_880 nm_400 um, DILAS, Tucson, AZ), collimating optics, and a digital micromirror display (DMD; DLP Light Commander; LOGIC PD, Carlsbad, CA) for generating optical patterns. The entire optical system was integrated with a slit lamp (Zeiss SL-120; Carl Zeiss, Thornwood, NY) for convenience of observing the illuminated retina in real time via a CCD camera (acA1300-60gmNIR; Basler, Ahrensburg, Germany).

To measure the stimulation threshold, NIR stimuli were applied at 2 Hz, with a pulse duration of 10 ms and peak irradiance ranging from 0.002 to 4.7 mW/mm^2^ on the retina. The light intensity was measured at the cornea and then scaled to the retinal irradiance by the ocular magnification squared, where the magnification was defined as the ratio between the sizes of the projected pattern on the retina and on the cornea. VEPs were recorded using the Espion E3 system (Diagnosys LLC, Lowell, MA) at a sampling rate of 2 kHz and averaged over 500 trials. The stimulation threshold was defined as the VEP amplitude exceeding the noise above the 95% confidence interval, as described previously [24].

## Results

### INL neurons

To assess the migration of the target bipolar cells and other inner retinal cells into honeycomb wells, 1 mm silicon devices (Fig. 1c-d) were implanted into the subretinal space of RCS rats (6 to 9 months old, n = 33) for 6 weeks. Each device comprised of four quadrants (Fig. 1c): flat, 20, 30 and 40 µm honeycombs to assess the effect of pixel size on retinal integration. The 25 µm height of the walls were chosen to accommodate the migration of second-order neurons (primarily BCs) and exclude the third-order neurons (ACs and RGCs) from the wells. Characterization of cellular integration with the implants was performed on reconstructed confocal acquisitions of the whole-mounted retina-honeycomb-sclera complex. Overview of the implant from the top of the honeycomb to the base of the well reveals a uniform fill by rod and cone bipolar immuno-labeled cells along with other non-labeled DAPI nuclei throughout the wells of different sizes (Fig. 2a, c, e). Cross-sectional (side) views through a randomly selected honeycomb row, projected from the middle of the wells to the sidewall, show migration to the base of the implants, while some BCs remain above the honeycomb walls. The fraction of the rod BCs, relative to its average number in the non-implanted control, inside the 25 µm tall walls of 20, 30 and 40 µm honeycombs was 0.64 ± 0.31, 0.77 ± 0.31, 0.53 ± 0.14, respectively (Fig. 2i). For cone BCs these fractions were 0.64 ± 0.19, 0.75 ± 0.15, 0.70 ± 0.17 respectively (Fig. 2j). Both bipolar cell types maintain their structural integrity, with unchanged stratification in the IPL.

**Figure 2.**
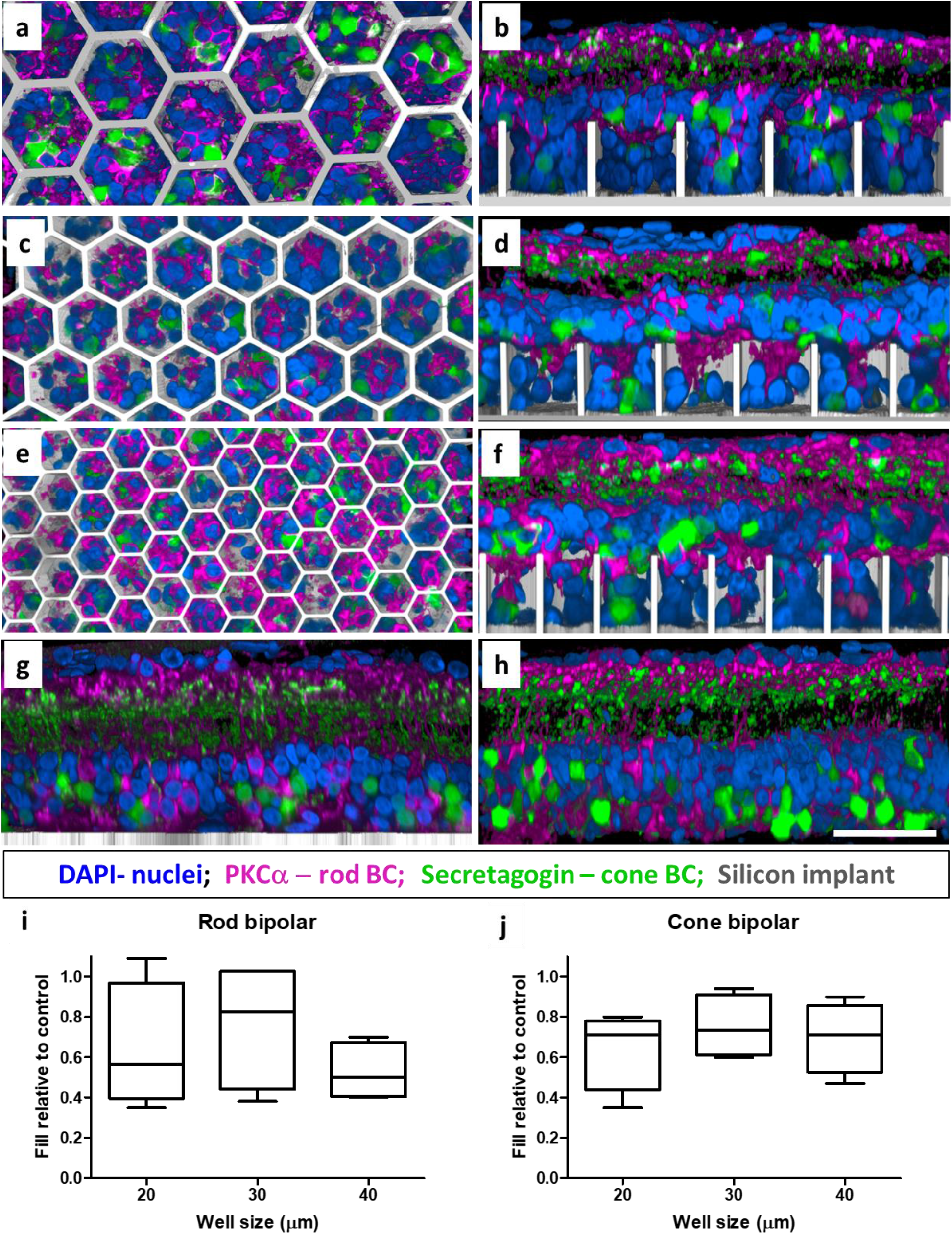
Confocal images of immuno-labelled rod (PKCα: magenta) and cone (secretagogin; green) bipolar cells. Top-down and side-view of the retina above and inside the honeycomb wells of 40 (a-b), 30 (c-d) and 20 (e-f) µm. DAPI (blue) labels the nuclei, and the implant is shown in grey. Scale bar = 50 µm. Bipolar cells maintain their structural integrity comparable to that with a flat implant (g) and a non-implanted RCS rat retina (h). Fraction of migrated rod (i) and cone (j) bipolar cells within the wells of 20, 30 and 40 µm, relative to non-implanted retina. Boxes extend from 25^th^ to 75^th^ percentile from the median line, and the whiskers - from smallest to largest value.

The other type of second-order neurons, horizontal cells, undergo dendritic and axonal degeneration, but remain in similar numbers after photoreceptor degeneration in RCS retina, compared to healthy retina [25]. Horizontal cells and their axons were observed close to the subretinal space in the non-implanted control (Fig. 3h) and interfacing with the flat implant control (Fig. 3g). In the presence of the 3-D implants, most horizontal cells migrated into the honeycombs: 0.85 ± 0.09, 0.70 ± 0.22 and 0.87 ± 0.17 in 20, 30 and 40 µm wells, respectively (Fig. 3i). However, after migration, the horizontal cells are now in the middle of the INL, close to the top of the wells (Fig. 3b, d, f). The top-down view shows cell bodies inside the wells, while their axons remain above the honeycomb walls (Fig. 3a, c, e; yellow arrows), as opposed to the natural morphology with their axons below the cell bodies (Fig. 3h).

**Figure 3.**
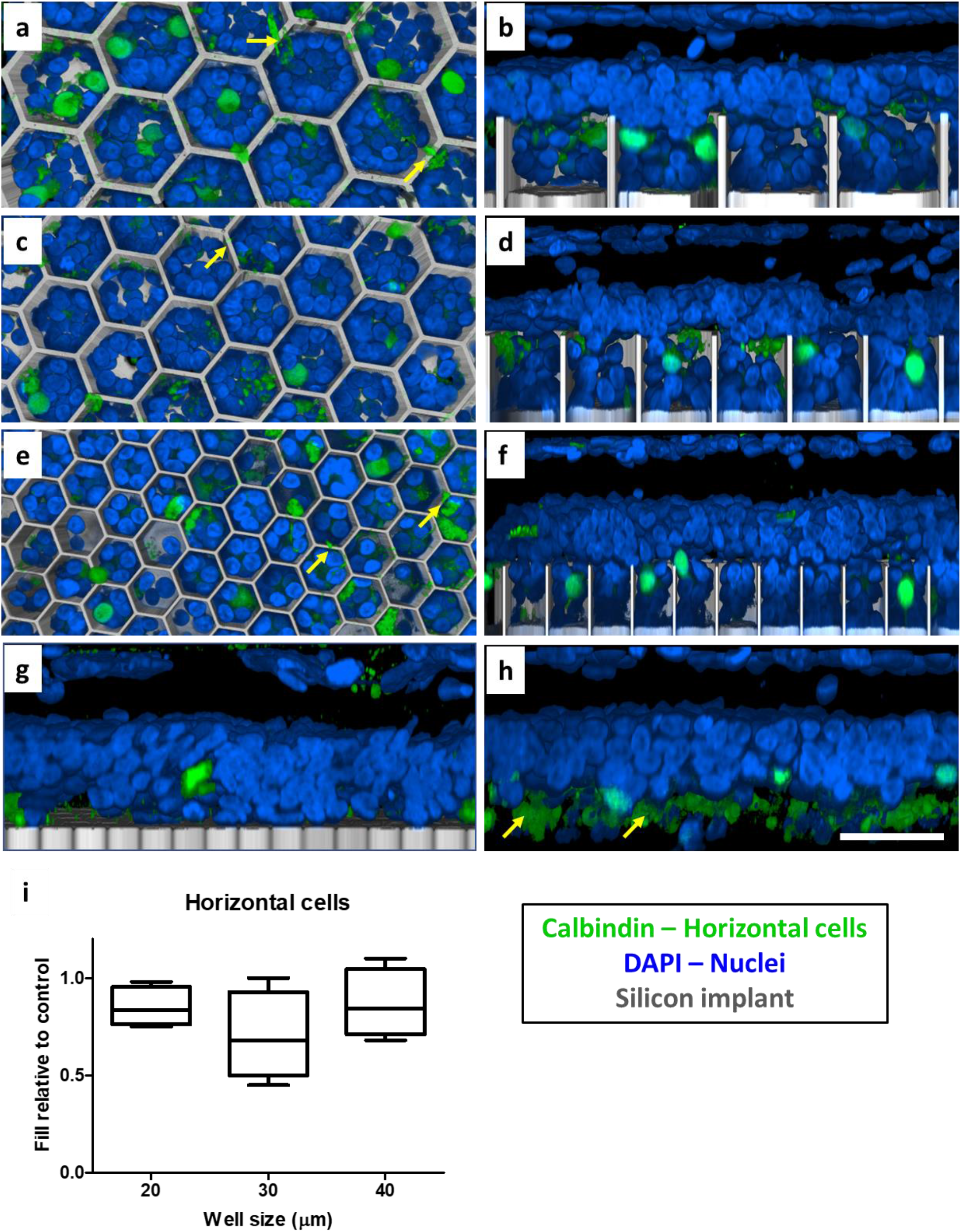
Confocal images of horizontal cells immuno-labelled with calbindin (green). DAPI (blue) label all other nuclei, and the implant is shown in grey. Scale bar is 50 µm. Top-down view (left column) and side view (right column) of the retina above and inside honeycomb wells of 40 (a-b), 30 (c-d) and 20 (e-f) µm. In the flat implant control (g) and non-implanted control (h), the cell bodies are above the axons (yellow arrows). In contrast, horizontal cells bodies migrate into wells, but their axons and dendrites (yellow arrows in a,c,e) remain above the walls. (i) Box extends from 25^th^ to 75^th^ percentile from the median line, and whiskers - from smallest to largest value.

The third-order neurons of the INL, amacrine cells, play a major role in signal transduction and modulation between bipolar and ganglion cells [26]. Therefore, direct electrical stimulation of the amacrine cells could lead to alteration of the natural signal processing. None of the immunolabeled subset of cholinergic starburst amacrine cells were observed inside the wells in the top-down view (Fig. 4a, c, e). Amacrine cell somas remain above the walls of all honeycomb sizes (Fig. 4b, d, f) away from the electric field, and preserve a similar IPL stratification as in the non-implanted control (Fig. 4h).

**Figure 4.**
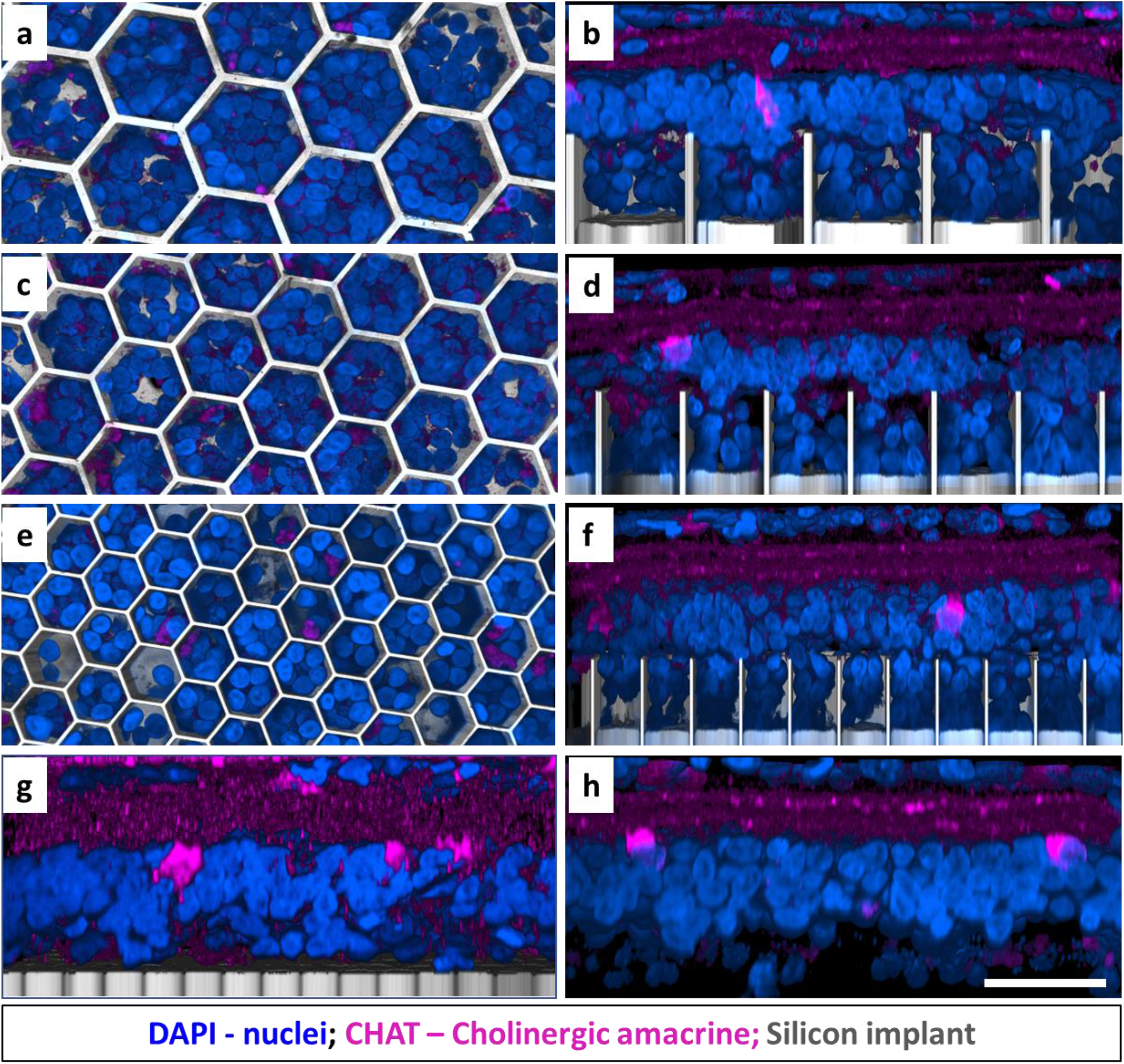
Amacrine cells immuno-labelled with choline acetyltransferase (CHAT: magenta). DAPI (blue) label the nuclei, and the implant is shown in grey. Scale bar is 50 µm. Top-down view (left column) and side view (right column) show the retina above and inside the honeycomb wells of 40 (a-b), 30 (c-d) and 20 (e-f) µm. Amacrine cells remain above the walls for all honeycomb sizes and retain their IPL stratification and cell body position in the INL, similar to the flat implant control (g) and non-implanted control (h).

### Inner retinal vasculature

An important consideration when dealing with retinal prosthesis is whether the device will allow normal oxygenation of migrated cells. Subretinal implants create a barrier between the choroidal supply and the retina. However, since the implants are inserted after a compete degeneration of photoreceptors, the choroidal supply is not necessary as the inner retina has its own vasculature. The inner retinal vasculature is grouped into the superficial vascular complexes (SVC; NFL to IPL) and the deep vascular complexes (DVC; IPL to OPL) [27]. The deep capillary plexus (DCP) of DVC comprises of vessels in the INL and OPL (subretinal space in degenerated RCS retina) as seen in Fig. 5g. Presence of a flat subretinal implant does not affect the DCP density or location, as compared to non-implanted area (Fig. 5h). With a 3-dimensional device, the DCP vessels rest on top of the walls (Fig. 5b, d, f) and do not migrate into the wells of any size studied. Nuclei (likely second-order neurons observed in Fig. 2-3) migrate past the vessels into the wells (Fig. 5a, c, e) and retain a healthy appearance 6-9 weeks after implantation, suggesting proper oxygenation and nutrients supply.

**Figure 5.**
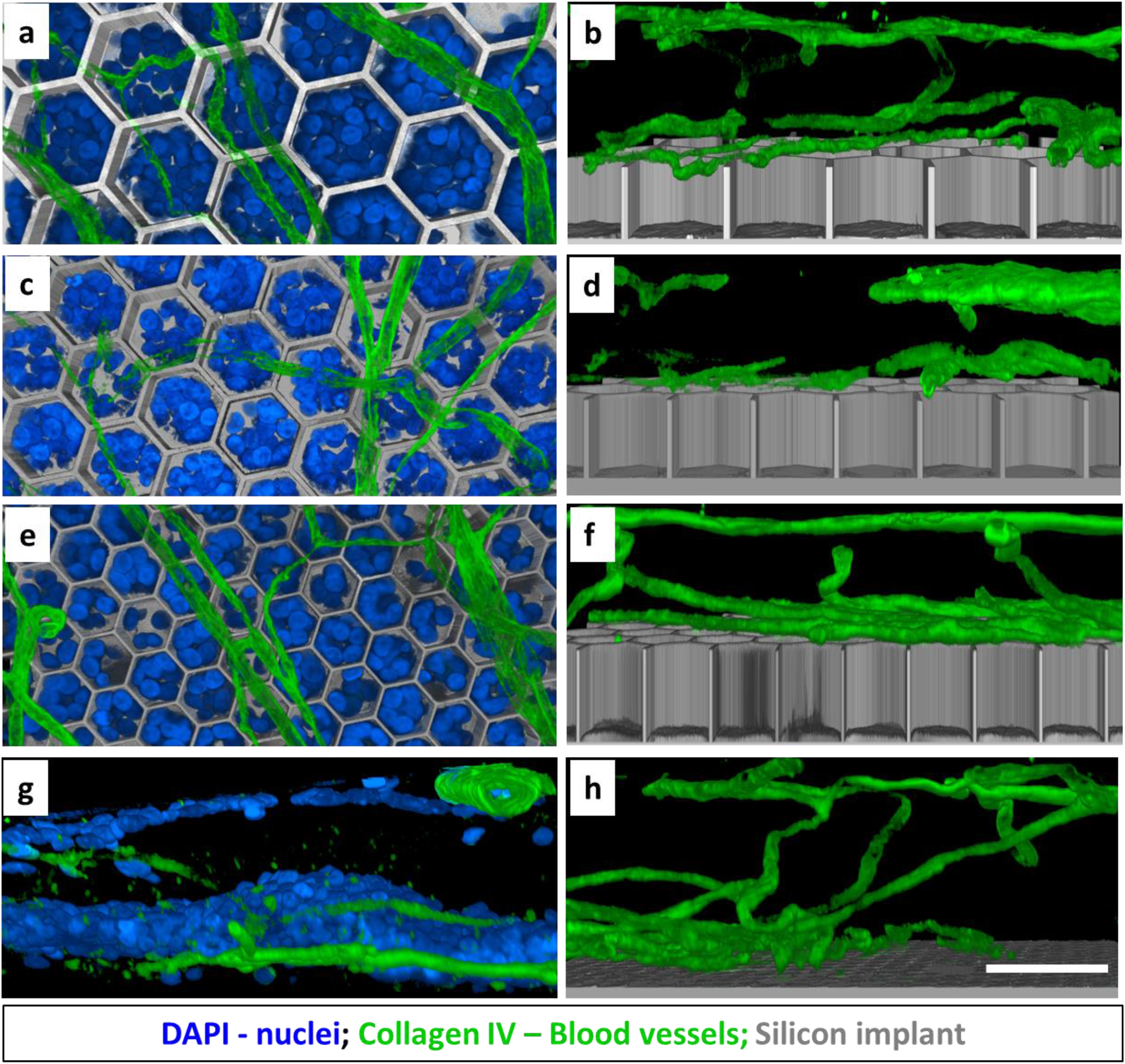
Retinal blood vessels immuno-labelled with collagen IV (green). DAPI (blue) label the nuclei, and the implant is shown in grey. Scale bar is 50 µm. Top-down view (left column) and side view (right column) show the vasculature above the honeycomb wells of 40 (a-b), 30 (c-d) and 20 (e-f) µm, while the retinal cells migrated around the vessels into the wells (a, c, e). The deep capillary plexus (DCP) interface with the subretinal space in a control retina (g) and with a flat implant (h).

### Retinal glial response

Müller glia span the entire thickness of the retina and ensheath all its neurons. Müller cells were immunolabelled by its cytoplasmic enzyme glutamine synthethase (GS), and Müller cell nuclei - by its transcription factor, SOX9. Migration of the Müller cell nuclei is known to happen after the retinal damage [28], similar to subretinal surgery for implantation of flat and 3-D arrays. In the non-implanted control, Müller cell nuclei are arranged orderly in the middle of the INL (Fig. 6h). After retinal integration with the honeycombs, some of the Müller cell processes and nuclei can be observed inside the wells in the top-down view (Fig. 6a, c, e). Most of the Müller nuclei migrate into the 30 µm and 40 µm wells: 0.73 ± 0.06 and 0.70 ± 0.08, respectively, but only 0.36 ± 0.12 into 20 µm wells (Fig. 6i) relative to the non-implanted control. The side views show some Müller cell bodies even reaching the bottom of the larger wells, but very shallow penetration into the 20 µm wells (Fig. 6b, d, f).

**Figure 6.**
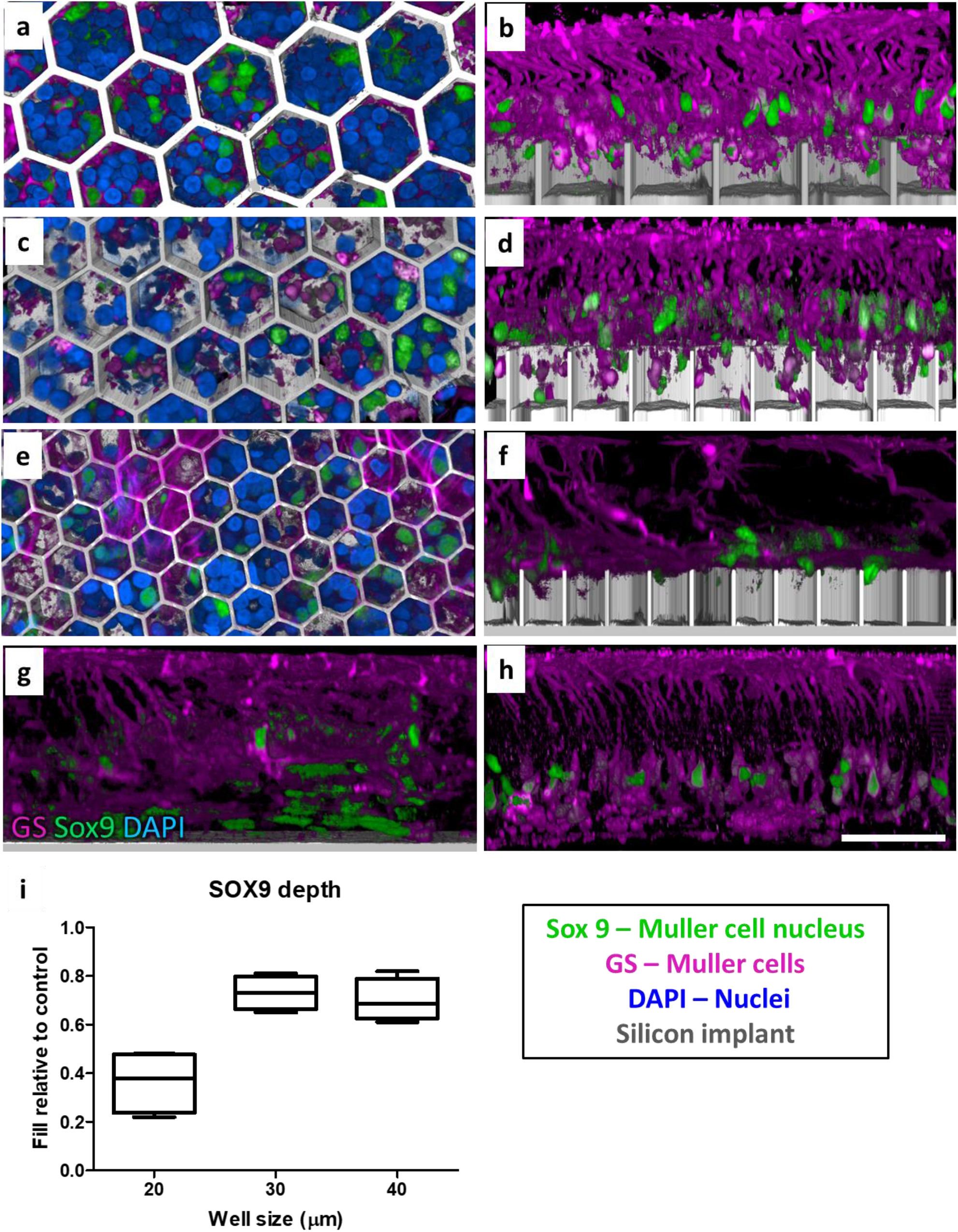
Muller cells (magenta) immuno-labelled with glutamine synthethase (GS) and Muller cell nuclei (green) labelled with SOX9. DAPI (blue) label other nuclei, and the implant is shown in grey. Scale bar is 50 µm. Top-down view (left column) and side view (right column) show the retina above and inside honeycomb wells of 40 (a-b), 30 (c-d) and 20 (e-f) µm. Muller cell processes and some of its nuclei migrate into the wells. (i) Depth of the SOX9 positive nuclei within the wells, compared to non-implanted control retina. Box extends from 25^th^ to 75^th^ percentile from the median line, and whiskers - from smallest to largest value.

Another consequence of a retinal insult is the Müller cell activation, which may lead to glial scar formation [28]. On a flat implant, glial scars may increase the distance and impedance between the active electrodes and the bipolar cells. This may be even more problematic with honeycomb implants as scar tissue could prevent migration of the bipolar cells into the wells and result in poor retinal stimulation. Müller cell activation was assessed by immunostaining the tissues with Glial Fibrillary Acidic Protein (GFAP), which is upregulated in Müller cells and astrocytes (Fig. 7). GFAP activation and clusters (indicative of a glial scar; yellow arrows) can be observed in the INL of the non-implanted RCS retina and flat implant control (Fig. 7g, h). Migration of the Müller cells into the wells increased with the size of the honeycombs: 0.43 ± 0.07, 0.60 ± 0.03, 0.71 ± 0.12 into 20 µm, 30 µm and 40 µm wells, relative to the position of GFAP staining in the INL in non-implanted controls. In contrast, the average penetration depth of all nuclei (DAPI staining in Fig. 7j) does not exhibit the well-width selectivity: 0.97 ± 0.08, 1.00 ± 0.11, 1.02 ± 0.06 of all INL nuclei, relative to the non-implanted control, migrate into 20 µm, 30 µm and 40 µm pixels, respectively.

**Figure 7.**
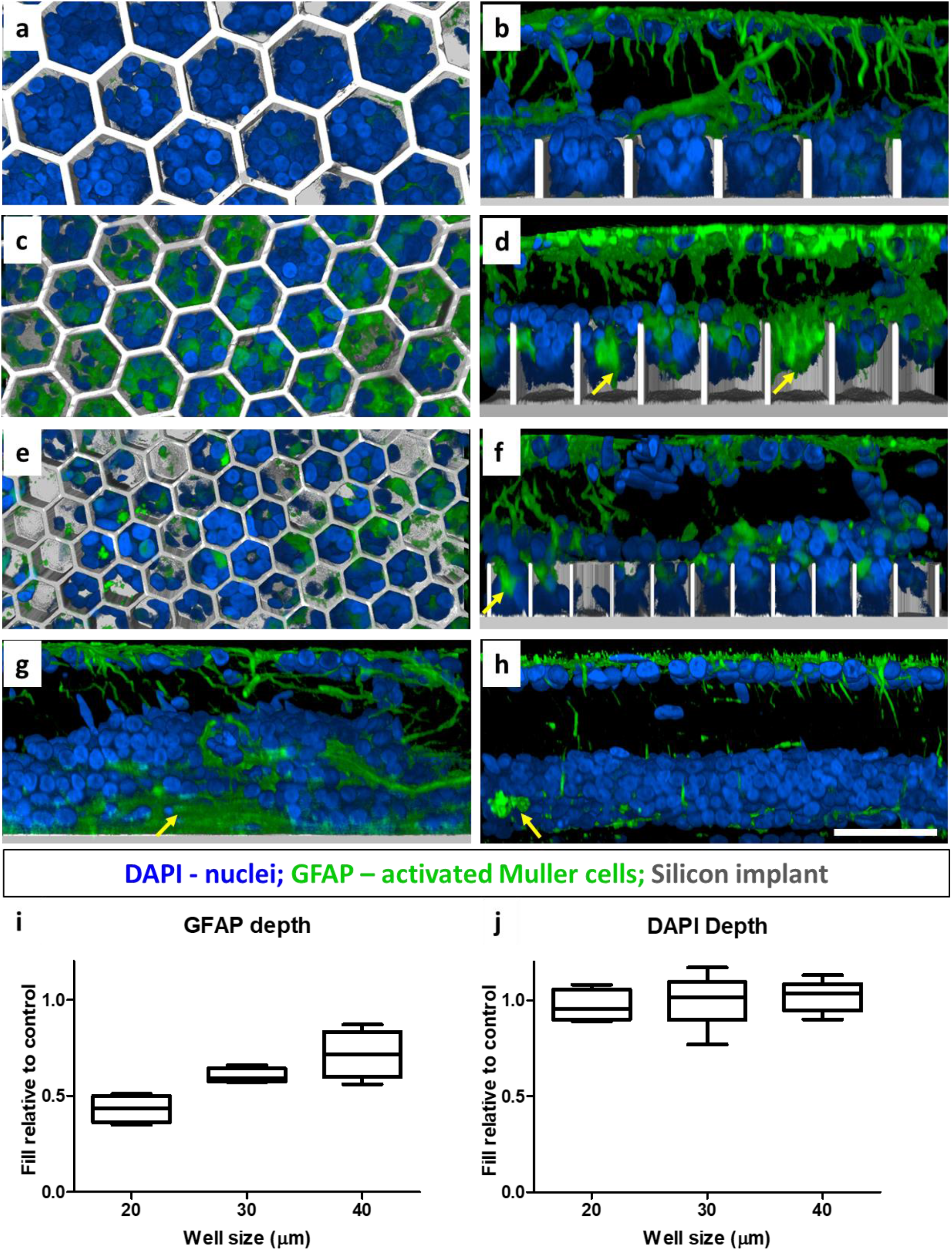
Muller cell activation marker GFAP (green). DAPI (blue) label the nuclei, and the implant is shown in grey. Scale bar is 50 µm. Top-down view (left column) and side view (right column) show the retina above and inside honeycomb wells of 40 (a-b), 30 (c-d) and 20 (e-f) µm. Yellow arrows point at large clusters of GFAP lacking the nuclei inside the wells, and also above the flat implant (g) and non-implanted control (h). Penetration of GFAP into wells (i) compared to DAPI nuclei penetration (j). Box extends from 25^th^ to 75^th^ percentile from the median line, and whiskers - from smallest to largest value.

Migration of neurons (Fig. 2 and 3) and all nuclei (Fig. 7j) deeper than Müller cell nucleus into the wells (specially in 20 µm wells) indicate that even in the event of glial scar formation (GFAP clusters and penetration in figure 7i), migration of the retinal neurons into honeycombs is not impeded.

### Retinal stimulation post migration

To assess whether the electrical excitability of the retina was affected by migration into the honeycombs, non-conducting vertical walls were formed in polymer structures on flat arrays with photovoltaic pixels, having a common return electrode only near the periphery of the array (monopolar configuration; Fig. 1e) [20]. The electric field in such wells is oriented vertically, similarly to that expected with an elevated return electrode on top of conductive walls. Such arrays were implanted into the subretinal space of RCS rats, temporal-dorsal to the optic nerve head (Fig. 8a). Surgical success was assessed using OCT immediately after the surgery, and migration was assessed six weeks after implantation. After surgery, the retina was separated from the implant by a thin layer of debris and fluid (Fig. 8 b,d). Six weeks after implantation, the INL is barely detectable by OCT above the honeycombs (Fig. 8c, e), but visible outside the implant, indicating the INL migration into the wells.

**Figure 8.**
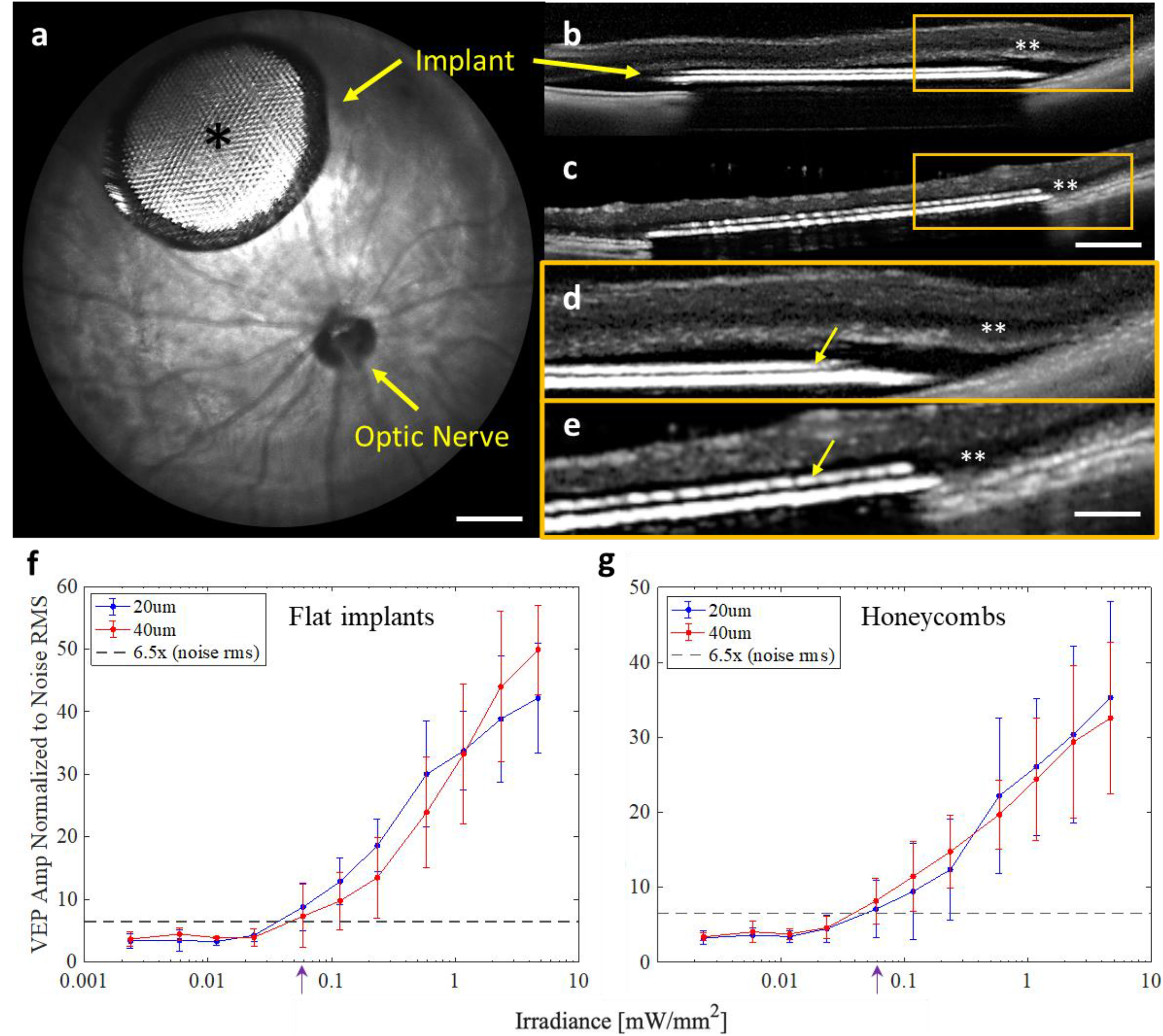
(a) Fundus of a rat eye with a subretinal honeycomb implant. The light area (*) of the implant is the photovoltaic pixels with honeycomb walls, surrounded by the darker return electrode. Scale bar = 500 µm (b) OCT image of the detached retina above the implant right after surgery and (c) 6 weeks later. Scale bar = 250 µm. Migration of the INL (**) into the honeycomb wells (yellow arrows) can be observed by comparing the higher magnification OCT at day 0 (d) and week 6 (e). Scale bar =100 µm. (f) The normalized VEP amplitude as a function of stimulus irradiance for the planar photovoltaic devices with 20 and 40 µm pixels. The black dashed line indicates the average noise level. The average threshold is 0.057 ± 0.029 mW/mm^2^ (g) The normalized VEP amplitude as a function of stimulus irradiance for 3D devices with polymer walls. The stimulation threshold is 0.064 ± 0.034 mW/mm^2^, similar to flat implants.

Visually evoked potentials (VEPs) were recorded via transcranial electrodes above the visual cortices, with NIR stimuli at 2 Hz, pulse duration of 10 ms and peak irradiance ranging from 0.002 to 4.7 mW/mm^2^ on the retina. The VEP was assessed with monopolar flat devices having 20 µm and 40 µm pixels and with 3-D printed walls on similar pixels for comparison (Fig. 8f,g). The stimulation thresholds with the 3D devices (0.064 ± 0.034 mW/mm^2^) closely matched that of full-field stimulation with the flat implants (0.057 ± 0.029 mW/mm^2^), where neighboring pixels combine to align the E-field vertically. This suggests that not only is the number of bipolar cells preserved post migration (Fig. 2), but so is the electrical excitability of the retina.

## Discussion

The recent clinical trials with flat photovoltaic implants having 100µm pixels (PRIMA, Pixium Vision SA, Paris, France) demonstrated restoration of central vision in AMD patients with acuity up to 20/438 [12], [13]. To further improve prosthetic visual acuity, pixel size should be decreased, while retaining sufficiently deep penetration of electric field into the INL, which is impossible with the flat bipolar pixel design. 3-D subretinal implants can address this limitation by decoupling the field penetration depth from the pixel width, thereby enabling smaller pixels. The elevated return electrode also aligns the electric field along the bipolar cells, decreasing the stimulation threshold, reducing the cross talk between neighboring pixels and providing better contrast [15]. In this study, we focus on the key aspects that the success of this technology is contingent upon, namely: a) migration of bipolar cells into the wells while retaining their electrical excitability, b) sufficient oxygenation and nutrient delivery inside the wells, c) minimal glial reactivity, and d) the wall height minimizing direct stimulation of the third-order neurons.

Retinal cell migration is a well-characterized phenomenon during ocular development [29]–[34]. Bidirectional movement of the newborn neurons is essential to stratification of the retinal tissue. In the degenerate retina, some level of migration also exists as part of the retinal remodeling process [4], [35], [36], [36]–[40]; Retinal response to laser damage of photoreceptors includes migration of the cone photoreceptors into the damage zone [41], bipolar cells rewiring to photoreceptors outside the damage zone [37], migration of the Müller cells nuclei [42], [36], [39], [28], and displacement of tertiary neurons [43]. Rapid and large-scale migration has been observed even ex-vivo, when a perforated membrane was placed on the photoreceptor side of the retina, while no migration was observed when such membrane was placed on the epiretinal side [19]. Similarly, rapid migration through a perforated subretinal membrane was observed with a degenerate retina in-vivo [19].

In this study, presence of the 3-D honeycomb implants induced a ‘de nouveau remodeling phase’, with migration of several cell types into the wells. Bipolar cells constituted the majority of migrated cells inside the honeycomb wells of all sizes, exceeding the estimated 8% of bipolar cells required for eliciting the VEP response [15]. The presence and stimulation of horizontal cells in the wells are unlikely to affect the retinal response since their normal synaptic connections with photoreceptors are missing, although random synaptic rewiring of horizontal cells in the degenerate retina [35], [37] cannot be excluded.

The optimal height of honeycomb walls is critical in allowing significant amount of the second-order neurons to migrate into the wells while providing their sufficient oxygenation and excluding the third-order neurons. Previously, we showed that 40µm deep cavities induce blood vessel migration and amacrine cell migration [18]. While blood vessels inside the wells may provide better oxygenation, the DCP are not numerous enough to migrate into every well within an implant. Furthermore, bending the laterally spanning vessels into cavities may induce damage to the vessel walls and compromise the retinal blood barrier. With 25µm high walls, the DCP and amacrine cells remain above the honeycomb walls while other INL cells migrate into the wells. The overall number of the cells in the INL is comparable to the controls, the cells retain a healthy morphology and more importantly, the retina remains electrically excitable, suggesting that proper diffusion of oxygen and nutrients is maintained over the induced 25µm-deep separation. Amacrine cells did not migrate into the wells and maintained their OPL stratification, contrary to the previous reports of the AC migration towards the GCL during retinal degeneration [36].

Müller cell activation and migration are a hallmark of retinal injury. During sustained retinal insult, Müller cell nuclear migration towards the injury site is thought to contribute to formation of a glial scar [44], [45]. In the degenerate retina, glial seal formation and progression is associated with the later stages of retinal remodeling [36]. In the presence of a 3-D array, the migration may be due to the surgical insult to the retina or a response to the same mediators that drive the other cell types. The fact that some neurons migrate deeper than the Müller cells nuclei and GFAP indicates that they are not impeded by the presence of a glial seal. While several mechanisms and drivers of retinal cell motility have been identified in the developing retina [29]–[31], migration mediators in the presence of our implants have not been characterized. If a bipolar cell-specific migration mediator can be identified, it could be leveraged to promote the bipolar cells migration, while other cell types might be halted.

The biological feasibility and compatibility of the honeycomb structures demonstrated in our study paves the way for decreasing the pixel size down to 20µm. Fabrication of the durable and active honeycomb devices, based on electroplating the conductive walls with return electrodes on top, is in progress. If successful in clinical trials, this technology may enable prosthetic vision with acuity exceeding 20/100, which would be very beneficial for many patients blinded by retinal degeneration.

---

**Supplemental Table 1.**
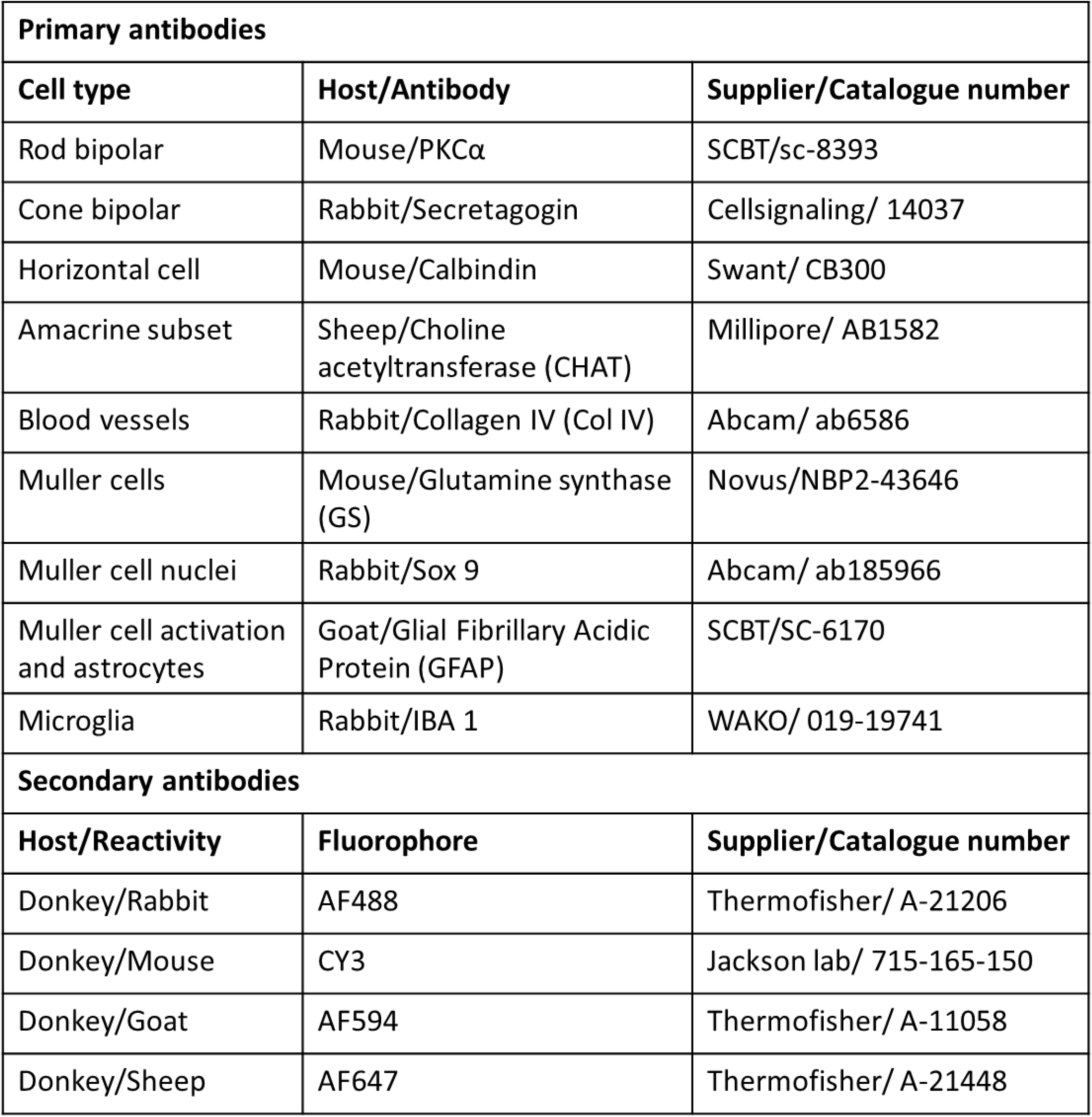
Primary and secondary antibodies used for immunohistochemistry.

## Acknowledgements

We would like to thank Ethan Rosenfeld from the Department of Physics at Stanford for his help with confocal image processing. Studies were supported by the National Institutes of Health (Grants R01-EY-027786, and P30-EY-026877), the Department of Defense (Grant W81XWH-19-1-0738), AFOSR (Grant FA9550-19-1-0402), Wu Tsai Institute of Neurosciences at Stanford, and unrestricted grant from the Research to Prevent Blindness. Photovoltaic arrays were fabricated at the Stanford Nano Shared Facilities (SNSF) and Stanford Nanofabrication Facility (SNF), which are supported by the National Science Foundation award ECCS1542152. K.M. was supported by a Royal Academy of Engineering Chair in Emerging Technology, UK.

## Notes

### Competing Interest Statement

DP-Patent & consultant (Pixium vision)
TK- Consultant (Pixium vision)

